# MutationalPatterns: comprehensive genome-wide analysis of mutational processes

**DOI:** 10.1101/071761

**Authors:** Francis Blokzijl, Roel Janssen, Ruben van Boxtel, Edwin Cuppen

## Abstract

Base substitution catalogs represent historical records of mutational processes that have been active in a system. Such processes can be distinguished by typical characteristics, like mutation type, sequence context, transcriptional and replicative strand bias, and distribution throughout the genome. MutationalPatterns is an R/Bioconductor package that characterizes this broad range of mutational patterns and potential relations with (epi-)genomic features. Furthermore, it offers an efficient method to quantify the contribution of known mutational signatures. Such analyses can be used to determine whether certain DNA repair mechanisms are perturbed and to further characterize the processes underlying known mutational signatures.

## Background

The genomic integrity of cells is constantly challenged by both endogenous and exogenous sources of DNA damage, such as UV-light and spontaneous reactions. Cells harbor a collection of DNA repair mechanisms to counteract these assaults. Not all lesions are, however, correctly repaired prior to replication, resulting in mutation incorporation into the genome [1]. Acquired mutations can have functional consequences and contribute to the development of diseases such as cancer and accelerate aging [2,3]. Knowledge on the causative mutational processes is therefore important for understanding disease etiology and could be valuable for future development of therapeutic strategies aimed at preventing or treating disease [4].

For example, the molecular defects of tumors are important determinants for treatment outcome, e.g., tumors that are deficient in homologous recombination (HR) are sensitive to compounds that increase the demand on HR, such as poly(ADP-ribose) polymerase (PARP) inhibitors [5]. There are, however, several ways through which a cellular pathway can be perturbed; in addition to germline and somatic mutations in known disease-associated genes, mutations in related genes or their regulatory elements can have a similar disruptive effect. Moreover, the dysregulation of genes can also occur through secondary mechanisms, such as DNA hypermethylation [6]. These various modes of deregulation make identification of molecular drivers at the individual gene level and stratification of patients towards the most optimal treatment challenging.

Alternatively, tumors can be stratified based on their altered activity of mutational processes. Each mutational process is thought to leave its own characteristic mark on the genome. For example, AID/APOBEC activity can specifically cause C>T and C>G substitutions at Tp<u>C</u>pA and Tp<u>C</u>pT sites (of which the underlined nucleotide is mutated) [7]. Thus, patterns of somatic mutations can serve as readouts of the mutational processes that have been active and can serve as proxies for the causal molecular perturbations in a tumor [8]. Generation and analysis of mutation catalogs of a patient’s tumor to guide diagnosis and treatment decision [8], is becoming increasingly conceivable for routine application, as whole-genome sequencing (WGS) costs are continuing to decline (https://www.genome.gov/sequencingcosts/).

In the past few years, large-scale analyses of human tumor genome data across different cancer types have revealed 30 recurrent base substitution patterns. These so-called “mutational signatures” are characterized by a specific contribution of 96 base substitution types with a certain sequence context [7]. Some mutational signatures are linked to specific processes through association with carcinogenic exposure, such as tobacco smoke [8,9], or the deficiency of DNA repair processes, such as nucleotide excision repair (NER) [10]. However, since multiple processes are typically disrupted in tumors, it is difficult to directly link a specific DNA repair and/or damage process to a signature based on genomic analyses of tumors. As a result, the etiology of the majority of mutational signatures identified in human cancers are currently unknown [11]. The underlying molecular mechanisms remain to be revealed in order to fully exploit mutational signature analysis for cancer diagnosis and treatment decision.

Until now, somatic mutation catalogs have been mainly determined for tumor samples, owing to their clonal nature. Recent and ongoing advances in single cell sequencing [12], extremely deep sequencing of clonal patches of healthy tissue [13,14] and clonal stem cell cultures [15] allow for the determination of somatic mutation catalogs of non-cancerous cells of various tissues. Furthermore, advances in gene-editing has enabled researchers to knock-out specific DNA repair mechanism and evaluate its effect on mutation accumulation [16]. For example, human adult stem cells in which the base excision repair (BER) protein NTHL1 was deleted using CRISPR-Cas9, showed a predominant increase of “signature 30” mutations [17], for which the underlying molecular mechanism was previously unknown. In a similar fashion, mutational signatures can be linked to specific sources of mutagenic stress, by studying their contribution in cells that are exposed to a specific carcinogen. To link damage or repair to previously known mutational signatures, it is essential to be able to quantify the activity of these mutational signatures in newly generated mutation catalogs.

In addition to mutational signatures, mutational strand asymmetries provide meaningful information on the underlying mutational processes. For example, transcriptional strand asymmetry arises in expressed genes through increased transcription-coupled NER (TC-NER) on the transcribed strand, and/or increased damage on the exposed untranscribed strand [18]. Decrease of this asymmetry potentially reveals a deficiency of TC-NER. Furthermore, replicative strand asymmetry can arise as a result of the different DNA polymerases that are used for the replication of the leading and lagging strand, which have distinct fidelities [18]. Increased replicative asymmetry may serve as a proxy for reduced proofreading capacities of polymerase ε (POLE) at the leading strand [19], or dysfunctional mismatch repair (MMR), which normally repairs most DNA polymerase mistakes [17,18]. These molecular defects have been observed in different human tumor types [18].

The distribution of mutations across the genome also provides valuable clues on the mutational mechanisms. For example, exposure to UV and alcohol increases the activity of error-prone DNA repair, involving translesion polymerase η (POLH), specifically at H3K36me3 chromatin in various cancer types. This effect does, however, not affect the overall mutation rate or spectrum. Rather, the carcinogenic effect is possibly a result of the differential targeting of mutations towards active genes, which are more likely to be consequential [20]. Analysis of the regional mutation rates in expressed genes and/or H3K36me3-associated regions is thus important to reveal this specific mutational mechanism. Finally, the distance between consecutive mutations can be evaluated to identify clustered mutagenesis called “kataegis”, a phenomenon associated with APOBEC overactivity [21], which has been shown to correlate with low responses to Tamoxifen [22,23].

Here, we describe MutationalPatterns, an R/Bioconductor package that allows researchers to easily evaluate and visualize a multitude of mutational patterns, such as type, sequence context, genomic distribution, association with genomic regions, transcriptional and replicative strand bias. Study of such characteristics enable deeper investigation of the molecular mechanisms underlying mutation accumulation. In addition, we have implemented a very efficient method to determine the contribution of known (e.g. COSMIC mutational signatures) or user-specified mutational signatures in individual samples. Using this method, it is possible to (1) estimate the contribution of known signatures in cells with (experimentally) altered DNA repair or damage and (2) to evaluate the activity of signatures in individual tumors. Taken together, MutationalPatterns is a versatile software package that facilitates the study of mutagenic agents and processes, the molecular dissection of existing mutational signatures, and the identification of molecular defects in individual tumors to improve diagnosis and treatment decision.

## Implementation

We implemented MutationalPatterns within the R/BioConductor platform [24], which is a widely used open-source software project for computational biology and bioinformatics. This platform provides easy integration with other R/BioConductor packages and workflows. All visualizations are generated with the powerful data-visualization package *ggplot2* [25], which can easily be adjusted to individual requirements with additional *ggplot2* commands. Moreover, publicly available genomic datasets can be retrieved using the *biomaRt* package [26] and used in the analyses, which allows exploration of a vast source of genomic annotation data from popular sources such as Ensembl (www.ensembl.org). In addition, in-house or publicly available experimental data such as RNAseq and ChIP-seq data, can be integrated.

### Data import and mutation types

Any set of base substitution calls, can be imported from a Variant Call Format (VCF) file and is represented as a *GRanges* object [27], a widely used data structure that allows very efficient computations including subsetting and overlapping with other genomic regions. MutationalPatterns reads VCF files in parallel, which reduces the time from *O(n)* to *O(n/c)*, where *n* is the number of VCF files, and *c* the number of cores available. All available reference genomes can be installed with the *BSGenome* package [28]. Once the data is imported, the sequence context of the base substitutions can be retrieved from the corresponding reference genome to construct a mutation matrix with counts for all 96 trinucleotide changes using “mut_matrix”. Subsequently, the 6 base substitution type spectrum can be plotted with “plot_spectrum”, which can be divided per sample group, such as tissue-type (Fig. 1A). Error bars indicate the standard deviation over the samples per group. For the C>T base substitutions, a distinction can be made between C>T at CpG sites and C>T at other sites, as deamination of methylated cytosines at CpG sites is a frequently active mutational process [7]. Moreover, a barplot with the 96 trinucleotide changes can be generated for each sample with “plot_96_profile”. Differences between two mutational profiles can be visualized using “plot_compare_profiles” (Fig. 2C), and the residual sum of squares (RSS) and cosine similarity values are indicated.

**Fig. 1.**
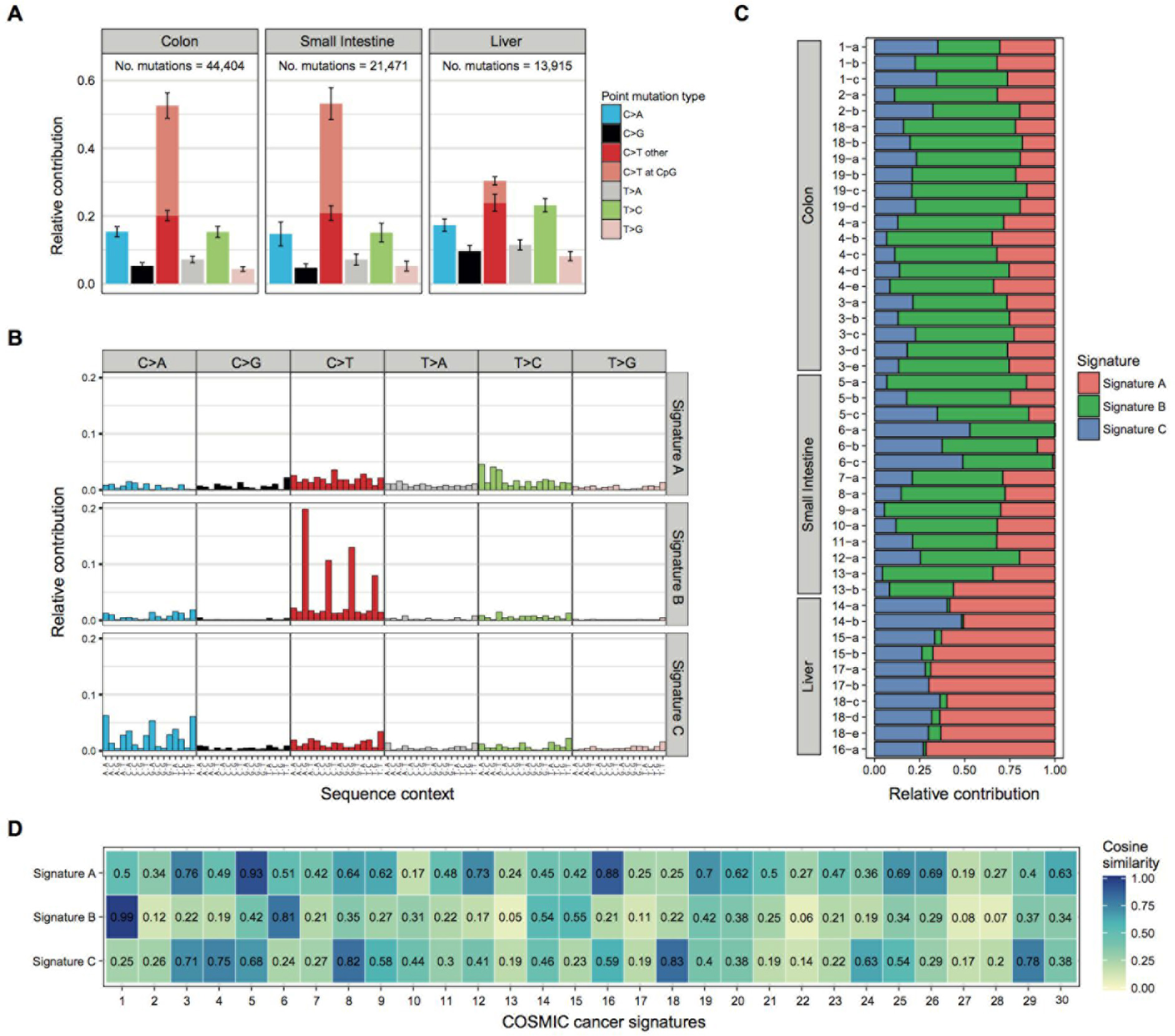
Characteristics of somatic mutations acquired in human ASCs of different tissues **a** Relative contribution of the indicated mutation types to the point mutation spectrum for each tissue type. Bars depict the mean relative contribution of each mutation type over all ASCs per tissue type and error bars indicate the standard deviation. The total number of somatic point mutations per tissue is indicated. **b** The relative contribution of each indicated trinucleotide change to the three mutational signatures that were identified by NMF analysis of the somatic mutation catalogs of the ASCs. **c** Absolute contribution of each mutational signature for each sample. Mutation loads between samples differ due to different ages of the donors. **d** Heatmap showing the cosine similarity of the mutational signatures in b. with the COSMIC signatures.

**Fig. 2.**
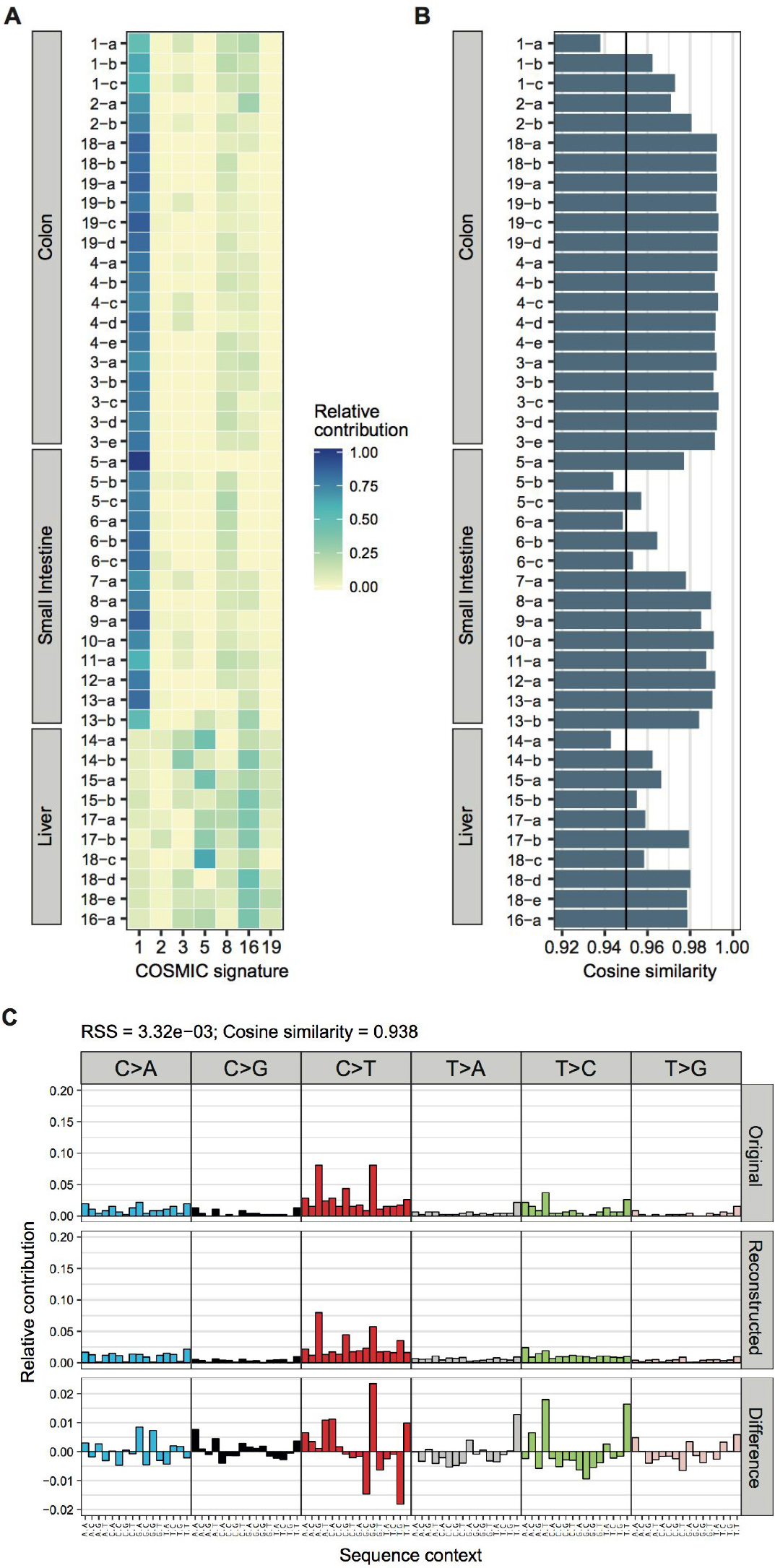
Using known mutational signatures to reconstruct mutational profiles of samples. **a.** The optimal relative contribution of COSMIC signatures to reconstruct the mutational profiles of the samples. The signatures with at least 10% contribution in at least one of the samples are plotted. **b.** The cosine similarity between the original mutational profile and the reconstructed mutational profile based on the optimal linear combination of all 30 COSMIC signatures. The line indicates the threshold of cosine similarity = 0.95. **c.** The relative contribution of each of the 96 trinucleotide changes to the original mutational profile (upper panel) and the reconstructed mutational profile (middle panel), and the difference between these profiles (lower panel) for the ASC with the lowest cosine similarity (*1-a*). The residual sum of squares (RSS) and the cosine similarity between the original and the reconstructed mutational profile are indicated.

### Mutational signatures

Mutational signatures can be extracted *de novo* from the mutation count matrix, which contains counts of all 96 trinucleotide changes in each sample, using non-negative matrix factorization (NMF) with “extract_signatures”. For this dimension reduction approach, the number of signatures is typically small compared to the number of samples in the mutation matrix. MutationalPatterns uses the implementation of R package *NMF* [29], which can also be used to estimate the optimal number of different mutational signatures that can be extracted from the data. Alternatively, novel probabilistic methods for identifying mutational signatures [30,31] can be used to extract signatures *de novo* and subsequent analyses can be carried out with MutationalPatterns. Mutational signatures can be visualized with “plot_96_profile”, and the contribution of each signature in each sample can be visualized in an absolute or relative barplot with “plot_contribution” (Fig. 1B), or a heatmap with “plot_contribution_heatmap” (Fig. 2A).

### Finding the contribution of known signatures in mutation catalogs

In addition to *de novo* signature extraction, the contribution of any set of signatures to the mutational profile of a sample can be quantified. This unique feature is specifically useful for mutational signature analyses of small cohorts or individual samples, as well as for relating own findings to known signatures and published findings. The non-negative linear combination of a set of user-specified mutational signatures that best reconstructs the mutation profile of a single sample, can be determined by minimizing the Euclidean norm of the residual, i.e.

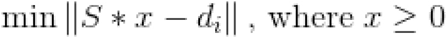

Here, *S* is the signature matrix, *x* the signature weight (contribution) vector and *d* the original 96 mutation count vector for sample *i*. This problem can be considered as a non-negative least-squares (NNLS) optimization problem, which is a constrained version of the least-squares problem where the weights are not allowed to become negative. The NNLS problem is well-studied, and a widely used algorithm for solving this problem is an active set method [32]. MutationalPatterns uses an R implementation of this algorithm from the *pracma* package [33] in “fit_to_signatures”.

### Mutational profile similarity

To determine the similarity α between two mutational profiles *A* and *B*, each defined as a non-negative vector with *n* mutation types, the cosine similarity is calculated:

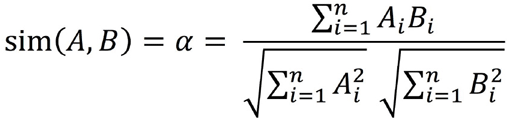

The cosine similarity can be calculated with “cos_sim” and has a value between 0 and 1. Two mutational profiles are identical when the cosine similarity is 1, and independent when the cosine similarity is 0. Because the cosine similarity evaluates the direction of the vectors and not the magnitude, it is not required to normalize the mutation profiles for the total number of mutations in a given sample. This similarity measure can be used for multiple MutationalPatterns analyses. Firstly, it can be used to calculate the similarity of *de novo* extracted signatures with previously described signatures using “cos_sim_matrix” (Fig. 1D). Secondly, the cosine similarity can be used to evaluate how well the reconstructed data - using either *de novo* or existing signatures - approximates the original data (Fig. 2B, C). Thirdly, the cosine similarity can be used to determine the similarity between the mutational profiles of samples and signatures using “cos_sim_matrix”, which can be visualized in a heatmap with “plot_cosine_heatmap” (Fig. 3). This heatmap does not represent signature weights to reconstruct mutation profiles such as found with “fit_to_signatures” (Fig. 2A), but rather reflects how well the mutational profile of a sample can be explained by each signature individually. As a result, each signature is given a cosine similarity value, and similar signatures have comparable values. The resulting plot visualizes the mutational similarity between the samples, while at the same time providing information on which signatures are likely active in the samples. Subsequently, the samples can be hierarchically clustered using the Euclidean distance between their cosine similarity values (Fig. 3).

**Fig. 3.**
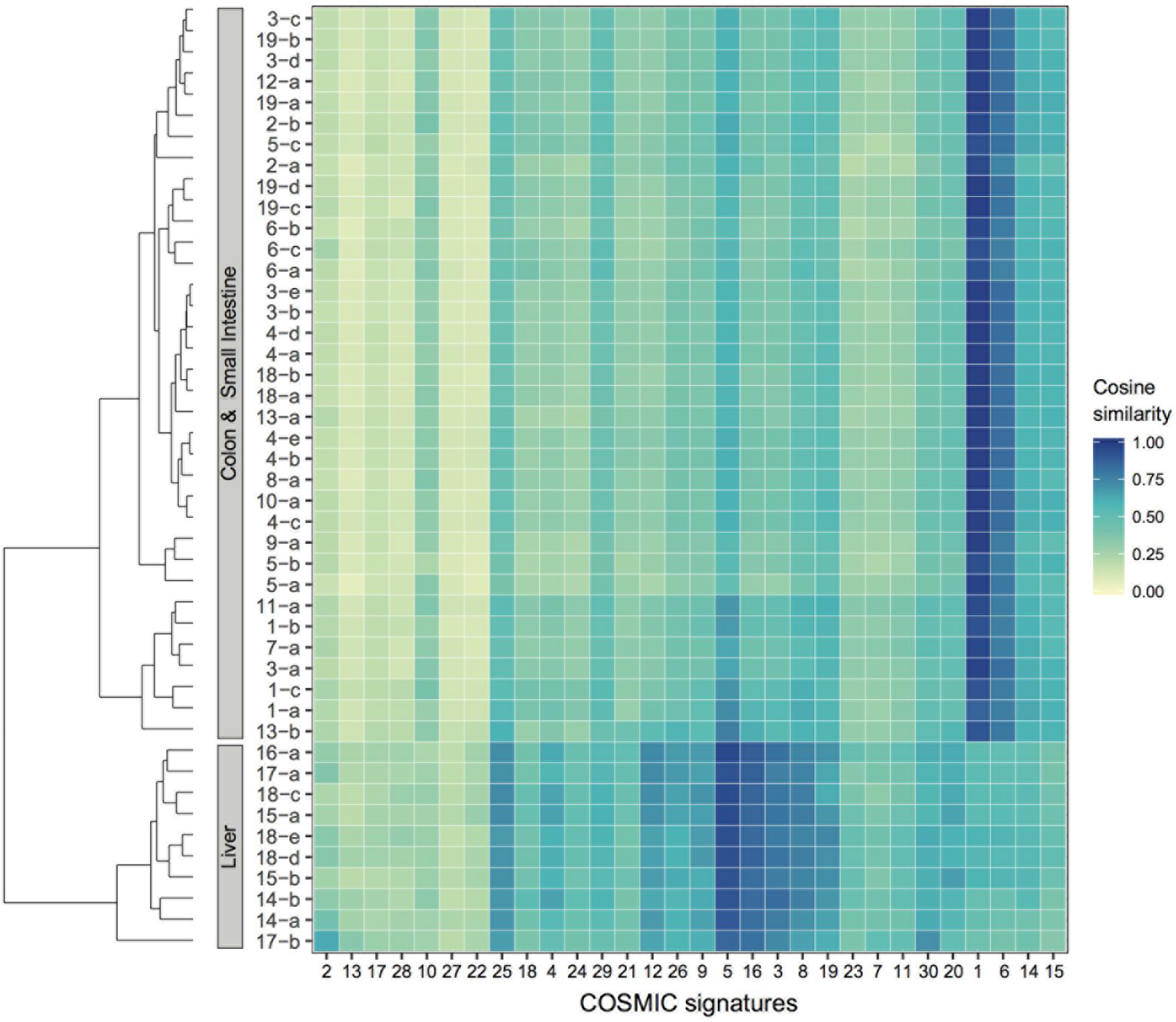
Heatmap with the cosine similarity between the mutational profile of each indicated sample and the 30 COSMIC signatures. The samples are hierarchically clustered (average linkage) using the Euclidean distance between the vectors of cosine similarities with the signatures. The signatures have been ordered according to hierarchical clustering (average linkage) using the cosine similarity between signatures, such that similar signatures are displayed close together.

### Mutational strand asymmetries

The involvement of transcription-coupled repair can be evaluated by testing for a transcriptional strand bias for the mutations that are located within gene bodies. While we cannot determine on which strand the original DNA damage occurred, we can regard the base substitutions from a reference frame of C>X or T>X changes (where X is any other base), and determine whether the mutated "C" or "T" base is located on the transcribed or non-transcribed strand. Since the gene definitions report the coding strand, which is untranscribed, base substitutions located on the same strand as the gene definitions are defined "untranscribed", and on the opposite strand as “transcribed”. Gene definitions for each reference genome can be retrieved from the UCSC genome browser [34] or BiomaRt [26] by loading a TxDb annotation package from Bioconductor. Subsequently, the transcriptional strand of all mutations within gene bodies can be determined with “mut_strand”.

The strand bias can be visualized for each sample with “plot_strand”, where the log2 ratio of the number of mutations on the transcribed and untranscribed strand is used as the effect size of the strand bias. A Poisson test can be performed to assess the statistical significance of the strand bias using “strand_bias_test” (Fig. 4C). In addition, the involvement of replication-associated mechanisms can be evaluated by testing for a mutational bias between the leading and lagging strand. The replication strand is dependent on the locations of replication origins from which DNA replication is fired. Replication timing is, however, dynamic and cell-type specific, which makes replication strand determination less straightforward. Replication timing profiles can be generated with Repli-Seq experiments [35]. Alternatively, replication timing dataset of human cell lines from the ENCODE project [36] are publicly available via the UCSC genome browser [34] and capture the conserved replication patterns. From replication timing profiles, the replication direction can be determined as described in [18]. Once the replication direction is defined, a strand asymmetry analysis can be performed using the same functions as for the transcription strand bias analysis.

**Fig. 4.**
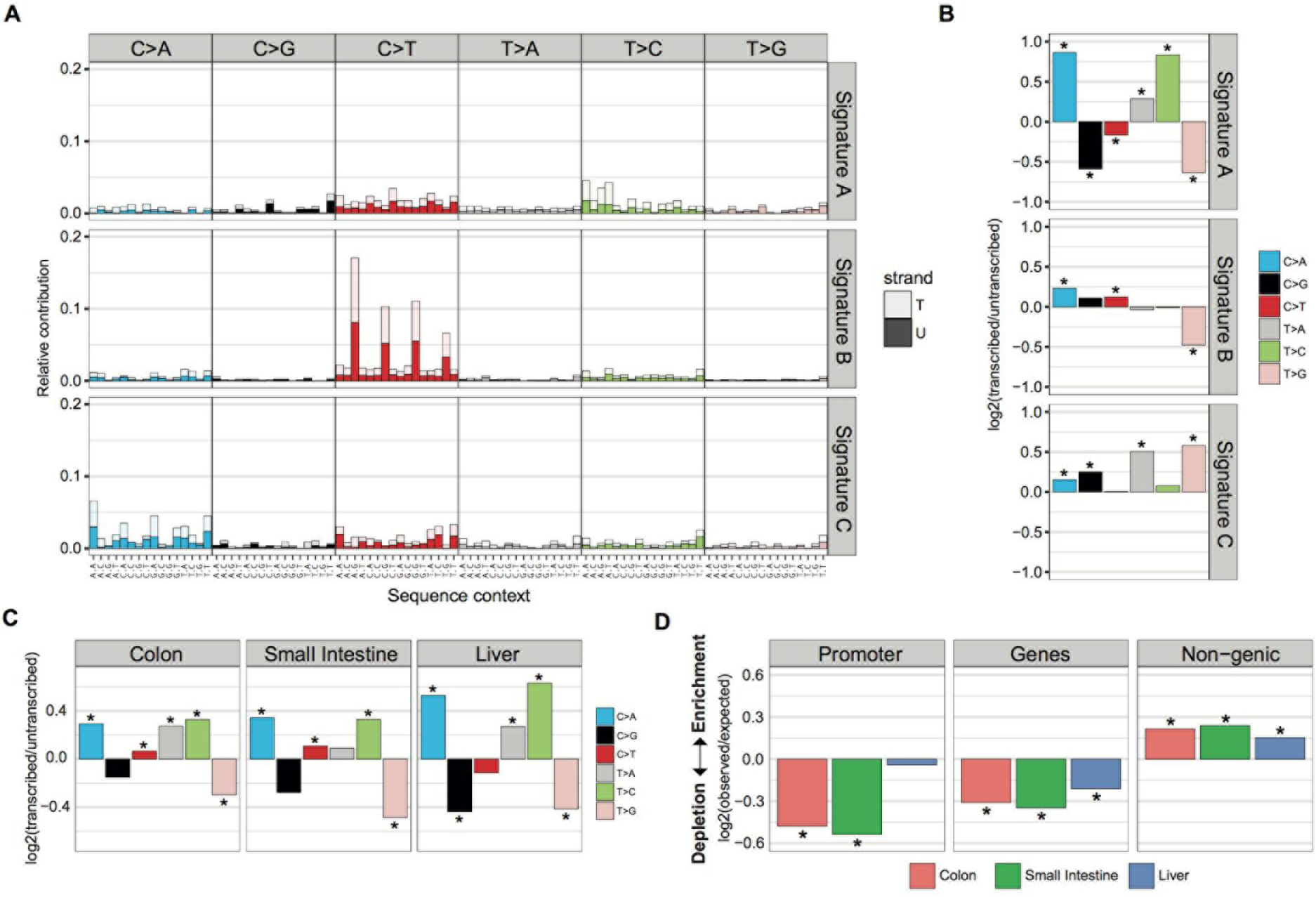
Transcriptional strand bias and genomic distribution. **a.** Mutational signatures with transcriptional strand information. The relative contribution of each trinucleotide change, subdivided into the fraction of trinucleotide changes present on the transcribed (T, light shades) and untranscribed strand (U, dark shades) **b.** Log2 ratio of the number of mutations on the transcribed and untranscribed strand per indicated base substitution for each signature depicted in a. The log2 ratio indicates the effect size of the bias and asterisks indicate significant transcriptional strand asymmetries (*P* < 0.05, two-sided binomial test). **C.** Log2 ratio of the number of mutations on the transcribed and untranscribed strand per indicated base substitution for each sample. Asterisks indicate significant transcriptional strand asymmetries (*P* < 0.05, two-sided Poisson test) **d.** Enrichment and depletion of somatic point mutations in the promoter regions, gene bodies, and intergenic genomic regions for all tissues. The log2 ratio of the number of observed and expected point mutations indicates the effect size of the enrichment or depletion in each region. Asterisks indicate indicate significant enrichments or depletions (*P* < 0.05, one-sided binomial test).

The transcriptional or replicative strand information can be included as an additional feature in the mutational signature analysis. Mutation count matrices with 192 features (96 trinucleotide changes * 2 transcriptional strands) can be created with “mut_matrix_stranded”. Subsequently, mutational signatures with 192 features can be extracted with “extract_signatures”, and their profile visualized as a stacked barplot with “plot_192_profile”. The effect size and the statistical significance of the strand bias of the signatures can be visualized with “plot_signature_strand_bias” (Fig. 4B).

### Genomic distribution

To determine whether base substitutions appear more or less frequently in specific genomic regions, the ratio of the observed and expected mutations in the genomic regions is determined with “genomic_distribution”. For this analysis, the chance of observing a mutation at one base is calculated as the total number of identified mutations, divided by the total number of bases in the genome that were surveyed in the sequencing experiment. Subsequently, the length of the genomic region that is surveyed is used to calculate the expected number of mutations in that genomic region. The “surveyed” bases are positions in the genome at which there are enough high quality reads to reliably call a mutation in that sample, and can be determined using the CallableLoci tool by GATK [37]. A list with GRanges of regions that were surveyed for each sample should be inputted in “genomic_distribution”. If a surveyed area would not be included in this analysis, it might result in e.g. a depletion of mutations in a certain genomic region that is solely a result from a low coverage in that region, and therefore does not represent an actual depletion of mutations.

Subsequently, the statistical significance of the enrichment or depletion is calculated with a one-sided binomial test with “enrichment_depletion_test”, and the enrichment or depletion can be visualized with “plot_enrichment_depletion” (Fig. 4). All genomic regions can be tested, as long as they are represented as GRanges objects [27]. The genomic regions can be based on experimental data or publicly available annotation data retrieved via e.g. BiomaRt [26], such as promoters, CTCF binding sites and transcription factor binding sites. Finally, a rainfall plot that visualizes the intermutation distance and mutation types can be made with “plot_rainfall” to identify localized hypermutation termed “kataegis”.

## Results and discussion

To provide a practical illustration of MutationalPatterns, we applied the various functionalities of the software package to somatic mutation catalogs of 45 human adult stem cells (ASCs) of different tissues [15]. The spectrum of base substitution types reveals a different mutational landscape for liver ASCs compared with intestinal ASCs (Fig. 1A), illustrating that this analysis can be used to detect gross differences in the activity of mutational processes. Deeper investigation into the processes can be achieved by performing a *de novo* extraction of mutational signatures using NMF.

We extracted three mutational signatures (Fig. 1B). Signature B, has a high contribution in intestinal ASCs specifically (Fig. 1C). The signature similarity analysis reveals that signature B is highly similar to COSMIC S1 (α = 0.99, Fig 1D), which is attributed to spontaneous deamination of methylated cytosines at CpG sites [7]. In liver ASCs, signature A shows the largest contribution, which was found to be similar to both S5 and S16 (α = 0.93 and 0.88 respectively, Fig. 1D). The underlying molecular mechanisms of these signatures are unknown, but both signatures are reported to have a transcriptional strand bias [11]. Consistently, transcriptional strand bias analysis of the mutation catalogs detects a strong bias for Signature A (Fig. 4A-B), confirming the likely involvement of transcription associated molecular mechanisms [18]. Lastly, signature C is most similar to COSMIC signature 18 (α = 0. 83, Fig. 1D), of which the etiology is currently unknown.

While the *de novo* signature extraction is a very powerful method for the identification of new signatures, it has several disadvantages. The analysis requires mutation sets with a large number of samples with diverse mutation spectra, as it relies on the dimensionality reduction method NMF. In order to evaluate the presence of the signatures in an additional sample, it must be added to the existing dataset and the complete analysis should be executed again. As a result, the input matrix will grow, and the runtime will increase with *O(n^3^)* where *n* is the number of samples, which makes this approach computationally demanding. Moreover, the extracted mutational signatures will slightly change every time a new sample is added.

Alternatively, the contribution of previously identified mutational signatures can be quantified in a single sample with “fit_to_signatures” feature of MutationalPatterns. Unlike NMF, this analysis is independent of other samples. Furthermore, the analysis is very fast with a runtime of approximately 0.1 seconds for 45 ASC samples (Supplemental Figure 1C), and is scalable with *O(n)* where *n* is the number of samples. This functionality can be used to study the activity of previously identified mutational signatures in cells with altered DNA damage or repair, which will help to uncover the molecular process underlying the mutational signature. Moreover, this analysis is suitable for clinical applications, as it allows for a fast per-patient analysis of the contribution of known signatures to their mutation profile.

By fitting the ASC mutational profiles to COSMIC signatures, we find comparable results as obtained with *de novo* signature extraction; the mutational landscape of intestinal ASCs is predominantly characterized by a high contribution of S1, and liver ASCs by both S5 and S16 (Fig. 2A). This result validates the ability of the “fit_to_signatures” function to identify active mutational processes by estimating the contribution of predefined signatures to mutation profiles. To test how reliable each mutational profile can be explained by the provided mutational signatures, the cosine similarity can be calculated between the original mutational profile and the mutational profile that is reconstructed using the determined optimal linear combination of mutational signatures (Fig. 2A). The mutational profiles of most ASCs can be reconstructed very well with the COSMIC signatures (mean α = 0.98, Fig. 2B), while some ASCs are not fully reconstructed (α < 0.95, Fig. 2B). This check is important, as a low similarity between the original and reconstructed profile indicates that the analyzed mutational profile cannot be fully explained by the provided signatures, which suggests that additional, unassessed mutational processes, might underlie the observed catalog of somatic mutations. Comparison of the original with the reconstructed mutational profile reveals which trinucleotide peaks cannot be reconstructed with the given signatures, and provides important leads on the missing mutational mechanisms active in the system studied (Fig. 2C).

Next, we determined the similarity between each mutational profile and each COSMIC signature, which reflects how well each mutational profile can be explained by each signature individually (Fig. 3). The advantage of this heatmap representation is that it shows in a glance the similarity in mutation profiles between samples, while at the same time providing information on which signatures are most prominent. Moreover, rather than choosing between similar signatures, it assigns a similarity score to each individual signature. Hierarchical clustering of the samples based on these profiles clearly separates the liver ASCs from the intestinal ASCs, while the colon and the small intestinal ASCs are not distinguishable by tissue-specific profiles (Fig. 3). This analysis demonstrates the utility of the MutationalPatterns package to detect and visualize sample groups with a similar activity of mutational processes.

Finally, we evaluated the enrichment and depletion of mutations in promoters, genes and non-genic regions. We downloaded these genomic annotations using biomaRt [26]. Intestinal ASCs show a depletion of mutations in promoter regions, whereas liver ASCs do not (Fig. 4C). This lack of depletion could be explained by binding of transcription factors to promoters, which can impair NER and result in increased rate of mutations at active promoters [38,39]. Furthermore, all ASC types show a depletion of mutations in genes and an enrichment in non-genic regions. This is expected, as genes are typically located in early-replicating genomic regions, where activity of MMR is known to be higher than in late-replicating regions [40]. In addition, expressed genomic regions may benefit from the presence of DNA damage repair through TC-NER and/or transcription domain-associated repair (DAR) [41,42]. The mutations in liver ASCs show the strongest transcriptional strand bias (Fig. 4C), indicating a high activity of TC-NER in these relatively quiescent cells. Nevertheless, the depletion in genes is larger in the intestinal ASCs compared with liver ASCs (Fig. 4C), which indicates that either replication-associated repair or DAR is more active in the highly proliferative intestinal ASCs. These results illustrate that the genomic distribution analysis provides important clues on the underlying mutational processes.

### Comparing methods

An overview of the functionalities of MutationalPatterns and related software tools can be found in Table 1.

**Table 1:**
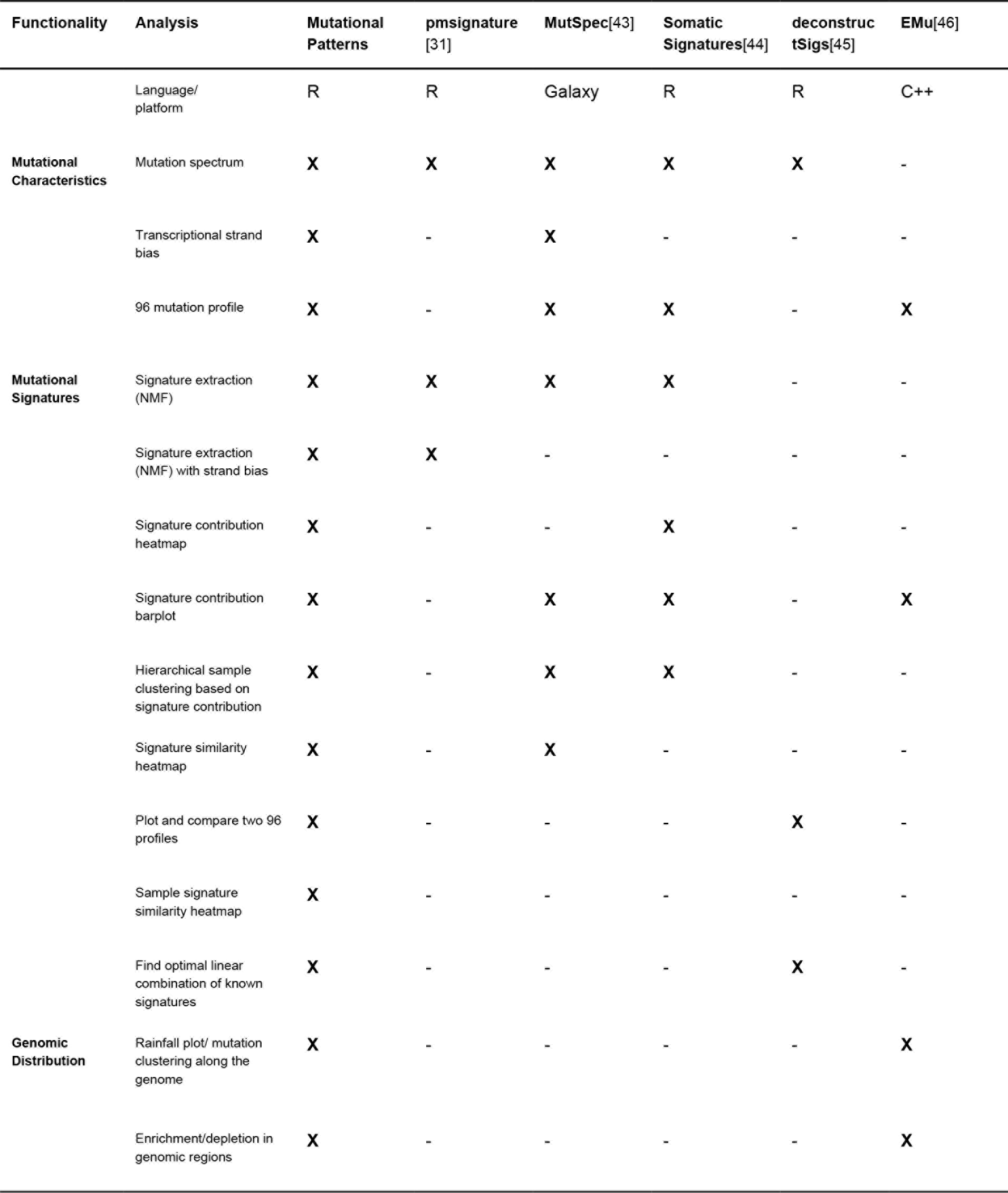
Feature overview and comparison with related software tools

An important advantage of MutationalPatterns over other available software tools is that it brings together many informative pattern analyses in a single package. Because MutationalPatterns is implemented within the R/Bioconductor platform, it integrates with common R genomic analysis workflows and allows easy association with publicly available annotation data. Moreover, MutationalPatterns can be used to easily generate publication-ready visualizations, while maintaining lay-out flexibility. The functionality to determine the activity of mutational processes through signature analyses in a single sample is an important feature. To date, only deconstructSigs provides this functionality, but has a different computational approach to this problem [45]. To determine the overlap in results and differences in functionality, we compared the performance of the “fit_to_signatures” function of MutationalPatterns with the “whichSignatures” of deconstructSigs. We used both functions to find the optimal linear combination of 30 COSMIC mutational signatures to reconstruct the somatic mutation profiles of the 45 human ASCs. The linear combinations of mutational signatures that were determined by these packages were highly similar (average Pearson correlation = 0.980, Supplementary Fig. 1a). To determine which package found the most optimal linear combination, we reconstructed the mutation profiles using the obtained linear combination of mutational signatures, and calculated their similarity with the original mutation profiles. The similarity was slightly higher for MutationalPatterns (mean α = 0.978) than for deconstructSigs (mean α = 0.977). Importantly, the MutationalPatterns analysis runtime is approximately 400 times faster compared with deconstructSigs.

## Conclusions

MutationalPatterns is a flexible and comprehensive R/Bioconductor package that allows researchers to rapidly assess a wide range of mutation characteristics in catalogs of somatic base substitutions. We showed that by analysing such patterns in concert, valuable clues on the molecular mechanisms underlying mutation accumulation can be revealed. MutationalPatterns allows researchers to generate publication-ready visualizations, which can be adapted easily to individual requirements.

In the past few years, mutational signature analyses have gained much interest, and some have been shown to have diagnostic value [8,10]. Since the etiology of most identified signatures is currently unknown, deeper investigation into the underlying molecular mechanisms will be essential to unfold signature analysis to its full potential. MutationalPatterns provides a very efficient method to determine the contribution of known mutational signatures in single samples, without requiring large data sets. This functionality will allow researchers to molecularly dissect well-established mutational signatures, by studying their contribution in cells with altered DNA damage or repair.

Finally, we anticipate that the ability to determine the activity of mutational signatures within individual patient samples has the potential to reveal molecular perturbations and thereby improve both diagnosis and treatment strategies. Furthermore, this analysis can facilitate biomarker discovery when the mutational signature activity is associated with treatment response. Taken together, we anticipate that MutationalPatterns will support fundamental research into mutational mechanisms, as well as enhance the knowledge that can be retrieved from individual patient sequencing data.

## Abbreviations

ASC: Adult Stem Cell; BER: Base excision repair; COSMIC: Catalog of Somatic Mutations in Cancer; HR: Homologous Recombination; NER: Nucleotide Excision Repair; NMF: Non-negative Matrix Factorization; NNLS: Non-Negative Least Constraints; MMR: Mismatch repair; PARP: Poly(Adp-Ribose) Polymerase; RSS: Residual Sum of Squares; TC-NER: Transcription-Coupled Nucleotide Excision Repair; VCF: Variant Call Format; WGS: Whole Genome Sequencing

## Declarations

### Ethics approval and consent to participate

Ethics approval as described in the article [15] from which we used the data that were analysed for this study.

### Consent for publication

Not applicable

### Availability and requirements

Project name: MutationalPatterns

Project homepage: https://github.com/UMCUGenetics/MutationalPatterns

Archived version: http://bioconductor.org/packages/release/bioc/html/MutationalPatterns.html

Operating system(s): Linux, Windows or MacOS

Programming language: R (version >= 3.4.0)

License: MIT

### Availability of data and code

The datasets supporting the conclusions of this article are included within the article [15] and are available at: https://wgs11.op.umcutrecht.nl/mutationalpatternsASCs/data/vcffiltered/ The code that can be used to reproduce all figures in this paper can be found at: https://github.com/UMCUGenetics/MutationalPatterns/blob/master/paper/figurespaper.R

### Competing interests

The authors declare that they have no competing interests.

### Funding

This work was financially supported by the NWO Gravitation Program Cancer Genomics.nl and the NWO/ZonMW Zenith project 93512003 to E.C.

### Authors' contributions

FB, RB and EC wrote the manuscript. FB developed and implemented the package. FB and RJ maintain the package. All authors read and approved the final manuscript.

## Acknowledgements

We thank the Bioconductor reviewers for their input on the R code.

---

**Supplementary fig. 1.**
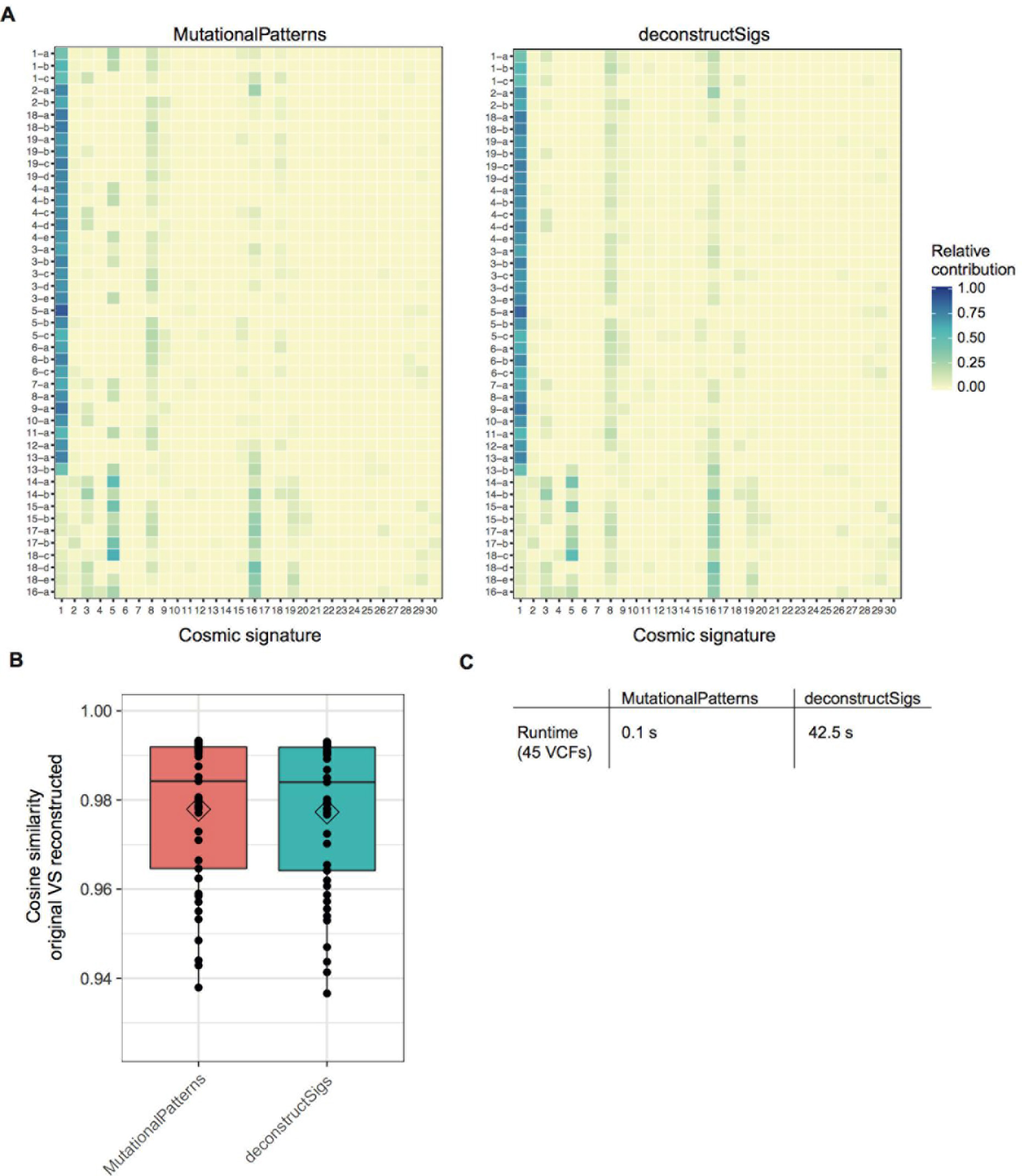
Comparison between the results of the functions of MutationalPatterns (fit_to_signatures) and deconstructSigs (whichSignatures) to find the optimal linear combination of a predefined set of mutational signatures to reconstruct a 96 mutation profile of a sample. **a.** Relative contribution of all 30 COSMIC signatures for each sample, as found by MutationalPatterns (left) and deconstructSigs (right). **b.** Cosine similarity between the original 96 mutational profile and the 96 mutational profile that can reconstructed with the linear combination of mutational signatures that was found by MutationalPatterns and deconstructSigs as depicted in panel a. **c.** The runtime (elapsed time) in seconds to find the optimal linear combination of mutational signatures for 45 somatic mutation catalogs for both packages; MutationalPatterns is approximately 400 times faster than deconstructSigs.

